# Mass graphs and their applications in top-down proteomics

**DOI:** 10.1101/031997

**Authors:** Qiang Kou, Si Wu, Nikola Tolić, Ljiljana Pasa-Tolić, Xiaowen Liu

## Abstract

Although proteomics has made rapid progress in the past decade, researchers are still in the early stage of exploring the world of complex proteoforms, which are protein products with various primary structure alterations resulting from gene mutations, alternative splicing, post-translational modifications, and other biological processes. Proteoform identification is essential to mapping proteoforms to their biological functions as well as discovering novel proteoforms and new protein functions. Top-down mass spectrometry is the method of choice for identifying complex proteoforms because it provides a “bird view” of intact proteoforms. The combinatorial explosion of possible proteoforms, which may result in billions of possible proteoforms for one protein, makes proteoform identification a challenging computational problem. Here we propose a new data structure, called the mass graph, for efficiently representing proteoforms. In addition, we design mass graph alignment algorithms for proteoform identification by top-down mass spectrometry. Experiments on a histone H4 mass spectrometry data set showed that the proposed methods outperformed MS-Align-E in identifying complex proteoforms.

## 1 Introduction

A proteoform is a protein product of a gene that may contain various primary structure alterations (PSAs) including: genetic variations, alternative splicing, and post-translational modifications (PTMs) [15]. The PSAs determine protein function in biological systems. For example, the combinatorial PTM patterns on histone proteins play a central role in epigenetic regulation [6,16]. Proteoform identification is essential to broadening our knowledge and deepening our understanding of proteoforms and their functions.

Despite the existence of various proteoforms, most protein sequence databases, such as Swiss-Prot [19], contain only one reference protein sequence for each gene or each transcript isoform. A complex proteoform may contain multiple PSAs compared with its corresponding reference sequence in the database (Fig. 1).

**Fig. 1.**
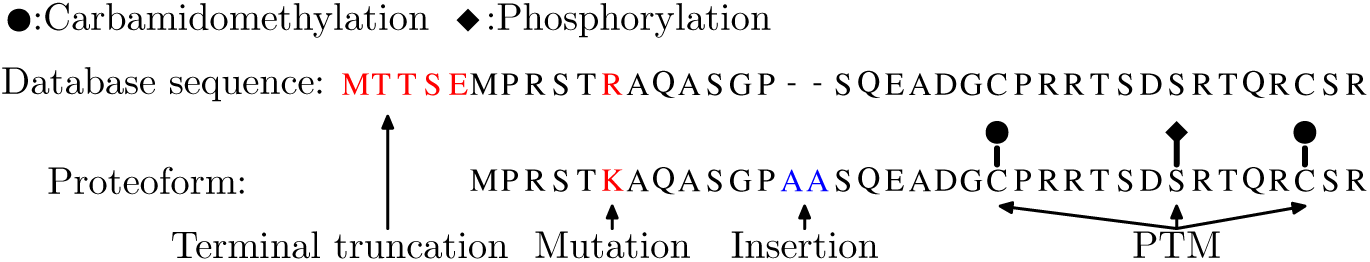
Comparison of a complex proteoform and its corresponding reference protein sequence in the database. The proteoform has an N-terminal truncation “MTTSE”, an amino acid mutation from “R” to “K”, an insertion of “AA”, one phosphorylated serine residue, and two modified cysteine residues with carbamidomethylation.

The differences between the target proteoform and its reference sequence make proteoform identification a challenging computational problem.

In proteoform identification, PSAs can be divided into several types: (a) sequence variations, such as mutations, insertions, and deletions; (b) fixed PTMs, which modify every instance of specific residues in the protein sequence; (c) variable PTMs, which may or may not modify specific residues in the protein sequence; (d) terminal truncations, which remove a prefix and/or a suffix of the protein sequence; and (e) unknown mass shifts of residues or subsequences, which are introduced by unknown PSAs. In Fig. 1, carbamidomethylation is a fixed PTM that modifies every cysteine residue; phosphorylation is a variable PTM that may modify serine, threonine, and tyrosine residues (only one serine residue is modified in the proteoform).

Top-down mass spectrometry (MS) has unique advantages in identifying proteoforms with multiple PSAs because it analyzes intact proteoforms rather than short peptides [4, 20, 22]. Recent developments in MS instrumentation and protein separation have paved the way for top-down MS analysis of complex proteoforms [2,17]. Fragment ion series in top-down tandem mass (MS/MS) spectra provide essential information to identify PSAs in proteoforms. Since top-down mass spectra are complex, they are often simplified by deconvolution algorithms [8, 10, 12] that convert fragment ion peaks into neutral fragment masses.

Let *S* be a spectrum of neutral fragment masses and *F* a proteoform with PSAs. Various scoring functions [14] for peptide-spectrum-matches in bottom up MS can be applied to measure the similarity of the proteoform-spectrum-match (*F*, *S*). Here we evaluate (*F, S*) using the *shared mass count score*, that counts the number of neutral masses in *S* explained by the theoretical neutral fragment masses of *F*.

Although the target protein of an MS/MS spectrum is generally unknown in proteome-wide studies, we can assume that the target complex proteoform is a product of a given known protein when purified proteins are analyzed. In this paper, we focus on the identification of proteoforms of known proteins with two types of PSAs: variable PTMs and terminal truncations. Fixed PTMs and amino acid mutations can be treated as special variable PTMs.

Let *P* be a reference sequence of the target proteoform and *Ω* a set of variable PTMs. We use *DB*(*P, Ω*) to represent the set of all proteoforms of *P* with Mass graphs and their applications variable PTMs in *Ω* and/or terminal truncations. Given a spectrum S, the proteoform identification problem is to find a proteoform *F ∈ DB*(*P, Q*) that maximizes the shared mass count score between *F* and *S*.

Extended proteoform databases and spectral alignment are the two main approaches for proteoform identification. ProSightPC [21] and MascotTD [9] use the first approach, in which spectra are searched against a sequence database of commonly observed proteoforms. However, the number of candidate proteoforms increases exponentially due to the combinatorial explosion of PTMs and truncations. For example, a protein containing 30 serine, threonine, or tyrosine residues has 2^30^ (about 1 billion) possible phosphorylated proteoforms. As a result, most uncommon proteoforms have to be excluded from the sequence database to keep its size manageable, limiting the ability to identify uncommon proteoforms.

Spectral alignment [7] is capable of identifying variable PTMs and unknown mass shifts since it finds a best scoring alignment between the spectrum and the reference sequence. However, existing alignment algorithms have their limitations. MS-Align+ [13] can identify proteoforms with at most two unknown mass shifts because it treats all PSAs as unknown mass shifts except for fixed PTMs and protein N-terminal PTMs. MS-Align-E [11] is capable of identifying proteoforms with variable PTMs, but not those with terminal truncations. MSPathFinder [1] is also capable of identifying variable PTMs, but truncations and unknown mass shifts are not considered in spectral alignment.

In this paper, we use mass graphs (Fig. 2) to efficiently represent proteoforms of a protein with variable PTMs and/or terminal truncations. In addition, mass graphs are capable of representing site specific variable PTMs. For example, we can specify that phosphorylation is a variable PTM for the second serine residue, but not other serine residues in Fig. 1. The idea of mass graphs is inspired by spectral networks [3] in bottom-up MS analyses and variant graphs [18] in proteogenomics studies. We transform the proteoform identification problem to the mass graph alignment problem and propose dynamic programming algorithms for a restricted version of the alignment problem.

**Fig. 2.**
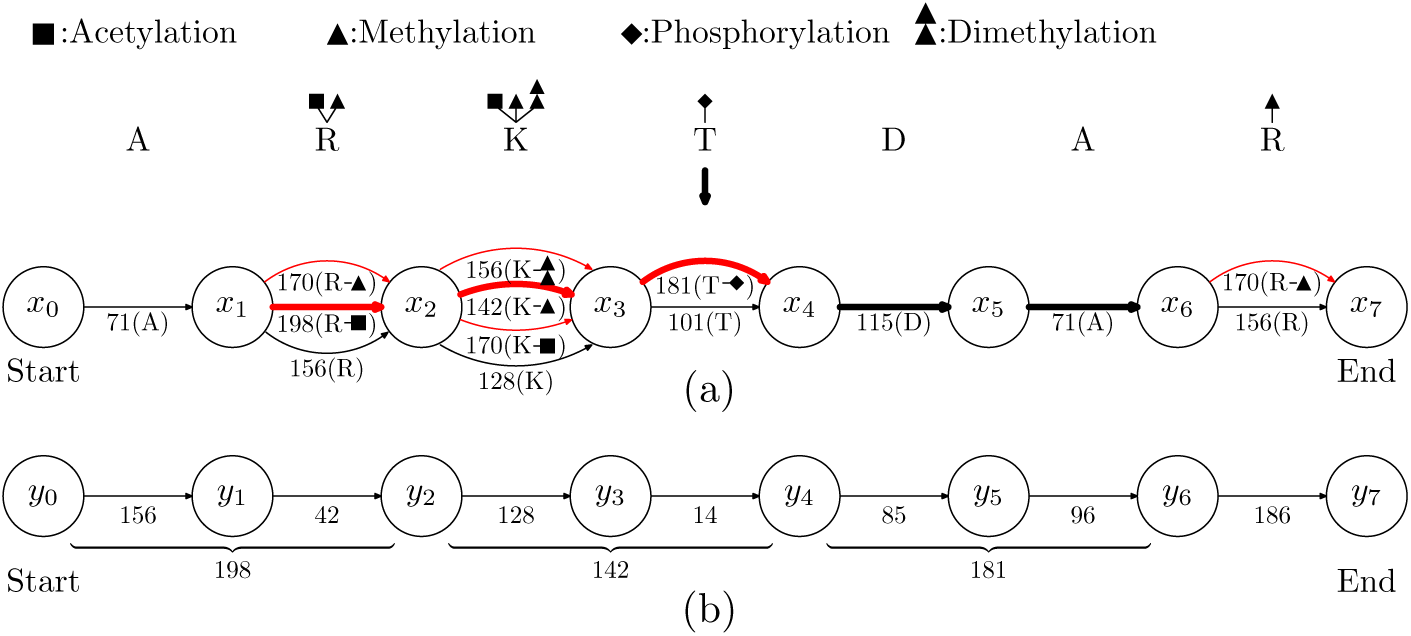
Construction of mass graphs. (a) An illustration of the construction of a proteoform mass graph from a protein ARKTDAR and four variable PTMs: acetylation on K and the first R; methylation on R and K, phosphorylation on T, and dimethylation on K. Each node corresponds to a peptide bond, or the N-or C-terminus of the protein; each edge corresponds to an amino acid residue (red edges correspond to modified amino acid residues). The weight of each edge is the mass of its corresponding unmodified or modified residue (a scaling factor 1 is used to convert weights to integers). (b) An illustration of the construction of a spectral mass graph from a prefix residue mass spectrum 0, 156, 198, 326, 340, 425, 521, 707. The spectrum is generated from a proteoform of RKTDA with an acetylation on the R, a methylation on the K, and a phosphorylation on the T. To simplify the mass graph, masses corresponding to proteoform suffixes (C-terminal fragment masses) are not shown. The full path from the start node *y*_0_ to the end node *y*_7_ is aligned with the bold path from node *x*_1_ to node *x*_6_ in (a). The path from *y*_0_ to *y*_6_ and the red bold path from *x*_1_ to *x*_4_ in (a) are consistent.

The proposed method were tested on a top-down MS/MS data set of the histone H4 protein. Experimental results showed that the proposed method outperformed MS-Align-E [11] in identifying complex proteoforms, especially those with terminal truncations.

## 2 Methods

Mass graphs are used to represent candidate proteoforms and top-down MS/MS spectra. Mass graphs representing proteoforms are called *proteoform mass graphs;* those representing MS/MS spectra *spectral mass graphs*. With the representation, we formulate the proteoform identification problem as the mass graph alignment problem and design dynamic programming algorithms for a restricted version of the problem.

### 2.1 Proteoform mass graphs

A proteoform mass graph is constructed from an unmodified protein sequence and its variable PTMs with three steps (Fig. 2(a)). (1) A node is added to the graph for each peptide bond of the protein. In addition, a start node and an end node are added for the N and C-termini of the protein, respectively. The *left node* of an amino acid is the one representing the peptide bond left of the amino acid. Specifically, the start node is the left node of the amino acid at the N-terminus. The *right node* of an amino acid is the one representing the peptide bond right of the amino acid. Specifically, the end node is the right node of the amino acid at the C-terminus. (2) For each amino acid in the protein, we add into the graph a directed black edge from its left node to its right node. The weight of the edge is the residue mass of the amino acid. (3) If an amino acid is a site of a variable PTM, we add into the graph a directed red edge from its left node to its right node. The weight of the edge is the residue mass of the amino acid with the PTM.

The locations of a PTM can be specified in a mass graph, thus reducing the number of candidate proteoforms. For example, the mass graph in Fig. 2(a) specifies that acetylation occurs on only the first arginine residue, not the second, in the protein. As a result, mass graphs are capable of representing amino acid mutations because a mutation can be treated as a variable PTM that modifies only the amino acid at the mutation site. To represent an amino acid with a fixed PTM, the weight of the black edge corresponding to the amino acid is assigned as the mass of the residue with the fixed PTM.

Each path in the graph represents a proteoform of the protein. A path from the start node to the end node is called a *full path* of the graph, representing a proteoform without terminal truncations. In the graph, the number of nodes is *O*(*n*), and the number of edges is *O*(*ln*), where *n* is the length of the protein sequence and *l* is the largest number of edges between two nodes.

### 2.2 Spectral mass graphs

Mass graphs are also used to represent top-down MS/MS spectra. In the preprocessing of spectra, peaks are converted into neutral monoisotopic masses of fragment ions by deconvolution algorithms [8,10,12]. Peak intensities are ignored to simplify the description of the methods. These monoisotopic masses are further converted to a list of candidate prefix residue masses, called a prefix residue mass spectrum [11]. The prefix residue mass spectrum of a collision-induced dissociation (CID) MS/MS spectrum is generated as follows: (1) Two masses 0 and *PrecMass – Water Mass* are added to the spectrum, where *PrecMass* is the precursor mass of the spectrum and *Water Mass* is the mass of a water molecule. (2) For each neutral monoisotopic mass *x* extracted from the spectrum, two masses *x* and *PrecMass − x* are added to the prefix residue mass spectrum. If the mass *x* corresponds to a proteoform suffix (prefix), then the mass *PrecMass − x* corresponds to a proteoform prefix (suffix).

A prefix residue mass spectrum with masses *a*_o_*, a*_1_*,…, a_n_* in the increasing order is converted into a spectral mass graph as follows (Fig. 2(b)). A node is added into the graph for each mass in the spectrum. The nodes for *a*_0_ = 0 and *a_n_* = *PrecMass − Water Mass* are labeled as the start and the end nodes, respectively. For each pair of neighboring masses *a_i_* and *a_i_*_+1_, for 0 ≤ *i ≤ n −* 1, a directed edge is added from the node of *a_i_* to that of *a_i_*_+1_, and the weight of the edge is *a_i−_*_1_ *− a_i_*. The spectral mass graph contains only one full path.

In the construction of mass graphs, the masses of all amino acids and PTMs are scaled and rounded to integers. A scaling constant 274.335215 is used to reduce the rounding error to 2.5 parts per million (ppm) [11]. Precursor masses and candidate prefix residue masses in highly accurate top-down mass spectra are discretized using the same method. As a result, all edge weights are integers in mass graphs.

### 2.3 The mass graph alignment problem

With the mass graph representation, the proteoform identification problem is transformed to an alignment problem between a proteoform mass graph and a spectral mass graph. The objective of the alignment problem is to find a path in the spectral mass graph and a path in the proteoform mass graph such that the similarity score between the two paths is maximized.

Let *A* be a path with *k* edges *e*_1_*, e*_2_*,…,e_k_*. The weight of the prefix *e*_1_*, e*_2_*,…, e_i_*, 1 ≤ *i* ≤ *k*, is called a prefix weight of *A*, denoted as *w_i_*. Specifically, *w*_0_ = 0 and *w_k_* is the weight of the whole path. The path *A* is also represented as a list of prefix weights *w*_0_*, w*_1_*,…,w_k_*. For example, the prefix weight list of the red bold path in Fig. 2(a) is 0, 198, 340, 521. Two paths are consistent if their weights are the same. For example, the red bold path in Fig. 2(a) and the path from *y*_0_ to *y*_6_ in Fig. 2(b) are consistent because they have the same weight 521.

We define the shared mass count score of two consistent paths *A* and *B* as the number of shared prefix weights in their prefix weight lists, denoted as Score(*A, B*). For example, the shared mass count score of the red bold path in Fig. 2(a) and the path from *y*_0_ to *y*_6_ in Fig. 2(b) is 4 because they share 4 prefix masses 0, 198, 340, and 521. If *A* and *B* are inconsistent, Score(*A, B*) = −∞.

#### Definition 1.

*Given a proteoform mass graph G and a spectral mass graph H, the mass graph alignment problem is to find a path A in G and a path B in H such that* Score(*A, B*) *is maximized*.

There are several variants of the mass graph alignment problem. In the local alignment problem (Definition 1), the two paths in the mass graphs are not required to be full paths (from the start to the end node). It can identify a sequence tag of the target proteoform as well as its matched masses in the spectrum. For example, the alignment between the red bold path in Fig. 2(a) and the path from *y*_0_ to *y*_6_ in Fig. 2(b) is a local alignment. The proteoform identification problem is transformed into the semi-global mass graph alignment problem in which the path *B* in the spectral mass graph is required to be the full path. If the path *A* is a full path, a proteoform without terminal truncations is identified. Otherwise, a truncated proteoform is reported. For example, the bold path (not a full path) from *x*_1_ to *x*_6_ in Fig. 2(a) is aligned with the full path in Fig. 2(b), corresponding to a truncated proteoform R[Acetylation] K[Methylation]T[Phosphorylation] DA. In the global alignment problem, both *A* and *B* are required to be full paths, that is, terminal truncations are forbidden.

We prove the mass graph alignment problem is NP-hard by reducing the subset sum problem [5] to the mass graph alignment problem. (See Appendix A.) Therefore, there are no polynomial time algorithms for this problem if P ≠ NP. In proteoform identification, we can reduce the search space by limiting the number of PTM sites in a proteoform. That is, only a small number of amino acids (sites) can be modified in a proteoform. This limitation gives rise to a variant of the mass graph alignment problem in which the number of red edges corresponding to modified amino acids is limited. The restricted mass graph alignment (RMGA) problem is defined as follows.

#### Definition 2.

*Given a proteoform mass graph G, a spectral mass graph H, and a number t, the restricted mass graph alignment problem is to find a path A in G and a path B in H such that A contains no more than t red edges and* Score(*A, B*) *is maximized*.

### 2.4 Consistent preceding node pairs

We use consistent preceding node pairs described below to solve the RMGA problem. In a mass graph, if there is a path from a node *u*_1_ to another node *u*_2_, we say *u*_1_ precedes *u*_2_. There may exist different paths from *u*_1_ to *u*_2_, each of which defines a distance that equals the weight of the path. Let *D*(*u*_1_*, u*_2_) denote the set of all the distinct distances defined by the paths from *u*_1_ to *u*_2_. The size of *D*(*u*_1_*, u*_2_) is much smaller than the number of paths from *u*_1_ to *u*_2_ when there are many duplicated distances introduced by consistent paths. For example, in Fig. 2(a), there are a total of 12 paths from *x*_1_ to *x*_3_, but *D*(*x*_1_*, x*_3_) contains only 7 distances {284, 298, 312, 326, 340, 354, 368}. When *u*_1_ is not a preceding node of *u*_2_, *D*(*u*_1_, *u*_2_) is an empty set.

Let *u*_1_, *u*_2_ be two nodes in *G* and let *v*_1_, *v*_2_ be two nodes in *H*. The node pair (*u*1*, v*1) is a consistent preceding node pair of the other node pair (*u*_2_*,v*_2_) if 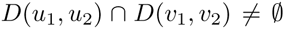, that is, there exist two consistent paths: one from *u*_1_ to *u*_2_, the other from *v*_1_ to *v*_2_. For example, the node pair (*x*_1_, *y*_0_) is a consistent preceding node pair of the node pair (*x*_3_, *y*_4_) in Fig. 2 because 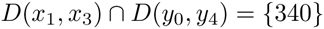.

#### Definition 3.

*Given a proteoform mass graph G and a spectral mass graph H, the consistent preceding node pair problem is to find all consistent preceding node pairs for every node pair* (*u, v*) *where u is in G and v is in H*.

Similar to the mass graph alignment problem, the consistent preceding node pair problem is NP-hard. We study a variant of the problem in which the number of red edges in a path in *G* is restricted. Let *D*(*u*_1_*,u*_2_*,r*) denote the set of distances defined by the paths from *u*_1_ to *u*_2_ that contain exactly *r* red edges, called an *r*-distance set. The node pair (*u*_1_, *v*_1_) is an *r*-consistent preceding node pair of the other node pair (*u*_2_*, v*_2_) if 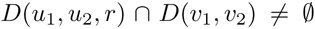. Next we describe algorithms for computing all *r*-distance sets of *G* and for finding *r*-consistent preceding node pairs based on *r*-distance sets.

*Algorithm for computing r-distance sets* Let *x*_0_*, x*_1_*, …,x_n_* be the nodes in the proteoform mass graph *G* in the topological order. We propose a dynamic programming algorithm (Fig. 3) for computing *D*(*x_i_, x_j_, r*) for 0 ≤ *i ≤ j ≤ n* and 0 ≤ *r* ≤ *t*. In the initialization (Steps 1 and 2), we set for each node *x_i_* in *G*

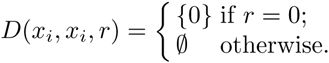

**Fig. 3.**
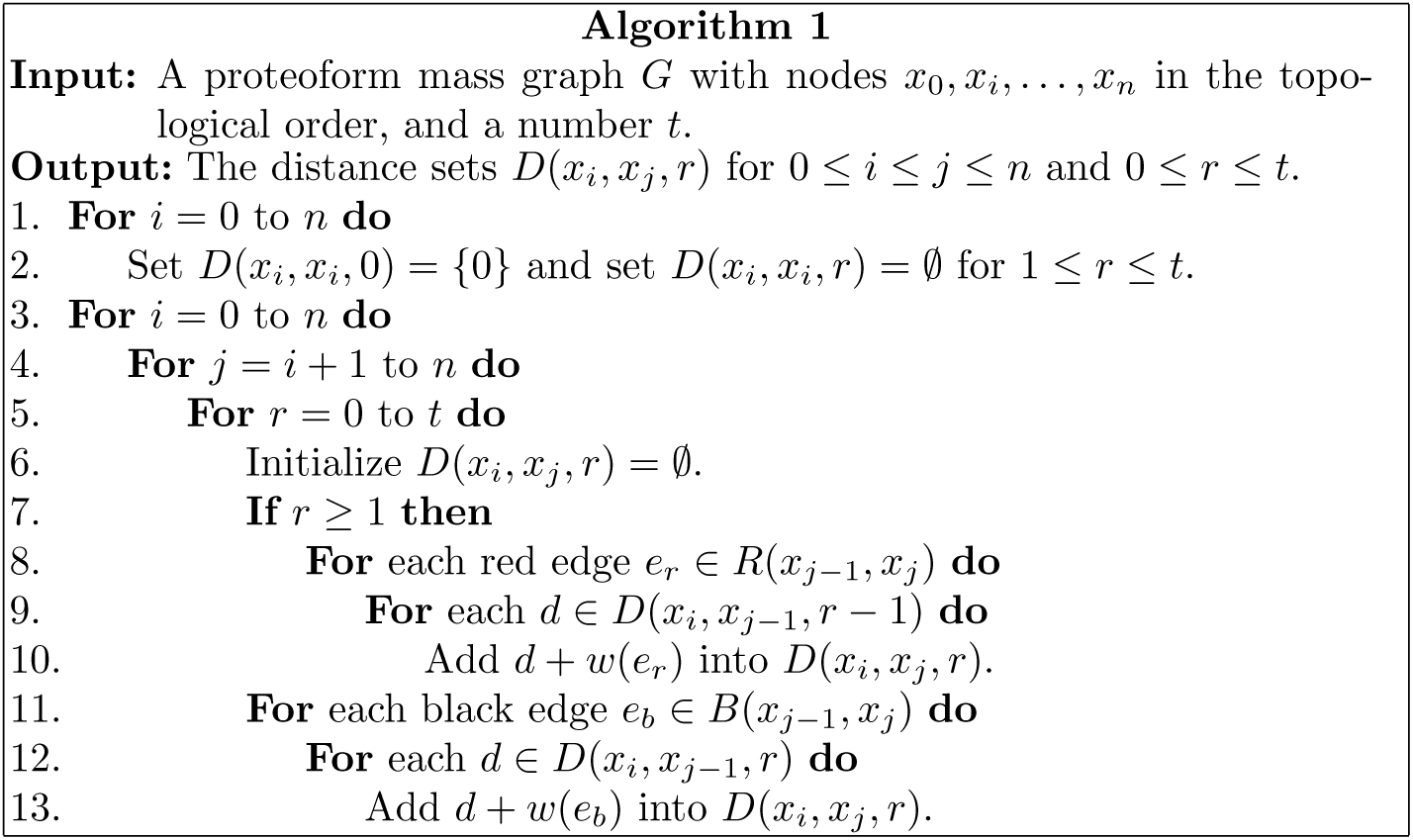
The algorithm for computing all the *r*-distance sets of a proteoform mass graph.

For 0 ≤ *i ≤ j ≤ n* and 0 ≤ *r ≤ t*, the set *D*(*x_i_, x_j_, r*) is computed based on the distances between *x_i_* and *x_j_*_−1_. Let *R*(*u*_1_*,u*_2_) (*B*(*u*_1_*,u*_2_)) be the set of all red (black) directed edges from a node *u*_1_ to another node *u*_2_. The weight of an edge *e* is denoted by *w*(*e*). For each red edge *e_r_* ∈ *R*(*x_j−_*_1_*, x_j_*) and each distance *d* ∈ *D*(*x_i_, x_j−_*_1_*, r −* 1), we add *d* + *w*(*e_r_*) into *D*(*x_i_, x_j_, r*) (Steps 7-10). For each black edge *e_b_* ∈ *B*(*x_j−_*_1_*, x_j_*) and each distance *d* ∈ *D*(*x_i_*, *x_j−_*_1_*, r*), we add *d* + *w*(*e_b_*) into *D*(*x_i_, x_j_, r*) (Steps 11-13).

The size of a distant set *D*(*x_i_*, *x_j_, r*) is *O*(*n^r^l^r^*), where *l* is the largest number of edges between two nodes in *G*. In the implementation, each distance set is stored in a sorted list, and Steps 12 and 13 are performed by merging two sorted lists with *O*(*n^r^l^r^*) steps. The time complexity of Steps 11-13 is *O*(*n^r^l^r+^*^1^). Similarly the number of operations of Steps 7-10 is also *O*(*n^r^l^r+^*^1^). The time complexity of Steps 5-13 is 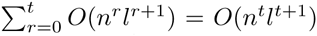, and the time complexity of the whole algorithm is *O*(*n^t^*^+2^*l^t^*^+1^).

The types of variable PTMs in proteoform identification are often limited. For example, only 5 types of PTMs were used in the experiments for the identification of proteoforms of the histone H4 protein. (See Section 3.2.) In this case, Algorithm 1 has a better time complexity. When a constant number *c* of PTM types are considered, the red edges in *G* can be divided into *c* types (variable PTMs). For example, the red edges in Fig. 2(a) are divided into four types based on their corresponding PTMs: acetylation, methylation, phosphorylation, and dimethylation. Each path in *G* has a *modification vector* [*z*_1_*, z*_2_*,… z_c_*] where *z_i_* is the number of red edges corresponding to the *i*th type of PTM. For example, the modification vector of the bold path in Fig. 2(a) is [1, 1, 1, 0]: one acetylation site, one methylation site, and one phosphorylation site. If two paths between two nodes have the same modification vector, they are consistent (their weights are the same) because their corresponding proteoforms have the same mass shifts introduced by PTMs. As a result, the size of a set *D*(*x_i_, x_j_, r*) is bounded by the number of different modification vectors satisfying that 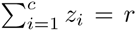, that is, the total number of red edges is *r*. The bound equals the number of ways to distribute *r* balls into *c* boxes, which is *O*(*r^c^*). Since the largest number of edges between two nodes *l ≤ c* + 1 is a constant, the time complexity of Steps 7-13 is *O*(*r^c^*). The number of operations in Steps 5-13 is 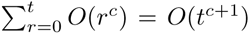, and the time complexity of the whole algorithm is *O*(*n*^2^*t^c^*^+1^).

We point out that the algorithm can be modified to find consistent preceding node pairs for general directed acyclic graphs (DAGs). A node *u*_1_ is an immediately preceding node of another node *u*_2_ if there is a directed edge from *u*_1_ to *u*_2_. In Steps 8-13 of the algorithm, we use *x_j_*_−1_, the only immediately preceding node of *x_j_*, to obtain *D*(*x_i_, x_j_, r*). For a general DAG, the node *x_j_* may have multiple immediately preceding nodes. In this case, all the immediately preceding nodes are used (similar to steps 8-13) to compute *D*(*x*_i_*, x_j_, r*).

*Finding r-consistent preceding node pairs* A node pair (*u*_1_, *u*_2_) in *G* and its *r*-distance set (*u*_1_, *u*_2_*, r*) = {*d*_1_*, d*_2_*,…,d_k_*} are represented by triplets < *u*_1_*,u*_2_*, d*_1_ *>,…,< u*_1_*, u*_2_*, d_k_ >*. For a given *r*, the triplets of all distance sets (●, ●*, r*) in *G* are merged and sorted based on the distance. Similarly, node pairs in *H* and their distances are also represented by a list of triplets sorted by the distance. The two sorted triplet lists are compared to find the *r*-consistent preceding node pairs for all node pairs (*u, v*) satisfying that *u* is *G* and *v* is in *H*. The time complexity of the step is *O*(*n^2^L*log(*nL*) + *m^2^* log *m* + *Z*), where *L* is the size of the largest *r*-distance set in *G*, *m* is the number of nodes in *H*, and *Z* is the total number of reported *r*-consistent node pairs.

Prefix residue masses in deconvoluted top-down MS/MS spectra may contain small errors introduced in measuring the *m/z* values of fragment ions. To address this problem, an error tolerance *ϵ* is used in finding *r*-consistent preceding node pairs. With the error tolerance, a triplet < *u*_1_*,u*_2_*,d_u_ >* from *G* matches a triplet < *v*_1_*, v*_2_*, d_v_ >* from *H* if |*d_u_* − *d_v_* | ≤ ϵ.

### 2.5 Algorithms for the RMGA problem

We present a dynamic programming algorithm (Appendix B) for the local RMGA problem. The algorithm can be modified to solve the semi-global and global RMGA problems. Let *x*_0_*, x*_1_*,…, x_n_* be the nodes in the proteoform mass graph *G* in the topological order, and let *y*_0_*, y*_1_*,…,y_m_* be the nodes in the spectral mass graph *H* in the topological order. We fill out a three dimensional table *T*(*i,j,k*) for 0 ≤ *i ≤ n*, 0 *≤ j ≤ m*, and 0 ≤ *k ≤ t*. The value *T*(*i, j, k*) is the highest shared mass count score among all consistent path pairs (*A, B*) such that *A* ends at *x_i_* and contains *k* red edges, and *B* ends at *y_j_*. Let *C*(*i, j, r*) be the set of all *r*-consistent preceding node pairs of (*x_i_, y_j_*). The values of *T*(*i, j, k*) are computed using the following function:

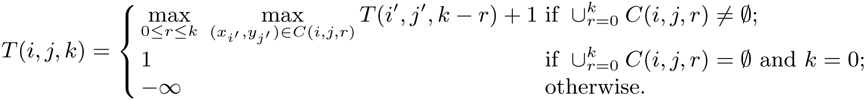

When (*x_i_, y_j_*) has no consistent preceding node pairs and *k* = 0, the value *T*(*i, j*, 0) is set as 1 because two empty paths have a shared prefix weight 0. After all values in the table *T*(*i, j, k*) are filled out, we find the largest one in the table and use backtracking to reconstruct a best scoring local alignment. The time complexity of the algorithm is *O*(*t^2^nmM*), where *M* the size of the largest set *C*(*i, j, r*). The algorithm can also be used for aligning two general DAGs.

The recurrence relation is slightly modified to solve the semi-global and global RMGA problems. For the semi-global alignment problem, we change the second line in previous recurrence relation to *T*(*i, j, k*) = 1 if 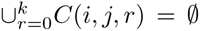 and *j* = *k* = 0, that is, *y_j_* is the start node. For the global alignment problem, we change the second line in the previous recurrence relation to *T*(*i, j, k*) = 1 if 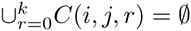 and *i* = *j* = *k* = 0, that is, both *x_i_* and *y_j_* are the start nodes.

## 3 Results

We implemented the proposed algorithms in C++ and tested it on a top-down MS/MS data set of the histone H4 proteins. All the experiments were performed on a desktop with an Intel Core i7-3770 Quad-Core 3.4 GHz CPU and 16 GB memory.

### 3.1 Data set

Core histones were separated by a 2D reversed-phase and hydrophilic interaction liquid chromatography (RP-HILIC) system of which the histone H4 protein was isolated in the first dimension. The protein separation system was coupled with an LTQ Orbitrap Velos (Thermo Scientific, Waltham, MA) to generate collision-induced dissociation (CID) and electron transfer dissociation (ETD) MS/MS spectra. A resolution of 60,000 was used for both MS and MS/MS spectra, and a total of 1, 626 CID and 1, 626 ETD spectra were acquired. More details of the MS experiment can be found in Ref [11].

### 3.2 Proteoform identification

We deconvoluted all the MS/MS spectra using MS-Deconv [12], and a window of 3 *m/z* was used for the deconvolution of precursor ions. Five common variable PTMs in the H4 histone protein (Table 1 in Appendix C) were included in the construction of the proteoform mass graph. For precursor masses, ±1 and ±2 Dalton (Da) errors were allowed, which may be introduced by the deconvolution algorithm. For a spectrum with a precursor mass *m*, we generated five candidate spectra with precursor masses *m* − 2, *m* − 1, *m*, *m* + 1, *m* + 2, respectively, and the spectrum with the best alignment result is reported. The error tolerance for fragment masses was set as *ϵ* = 0.1 Da and the largest number of red edges (PTMs) *t* was set as 10. By aligning the spectra against the proteoform mass graph, the algorithm for the semi-global RMGA problem identified a total of 1, 183 proteoform-spectrum-matches with at least 10 matched fragment ions, including 999 matches with at least 20 matched fragment ions (Fig. 5(a) in Appendix D). Of the 1, 183 matches, 578 contain more than 3 PTM sites (Fig. 5(b) in Appendix D).

The running time of the proposed approach was about 800 minutes. The running time depends on the sizes of the *r*-distance sets and the numbers of *r*-consistent preceding node pairs reported from the proteoform and spectral mass graphs. For the histone H4 protein with the five variable PTMs, the size of the largest r-distant set was 891. For each spectral mass graph, we count the total number *N* of the consistent preceding node pairs used in the mass graph alignment algorithm, that is, 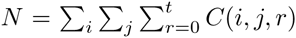. The average value of *N* for all the 3, 252 spectra was 1.27 × 10^7^, and the maximum value of *N* was 1.73 × 10^8^.

### 3.3 Comparison with MS-Align-E

We compared the performance of the proposed algorithms and MS-Align-E [11]. For MS-Align-E, the error tolerance for fragment masses was set as 15 ppm and all the other parameters were set as the same as the mass graph alignment method. MS-Align-E identified 1, 031 proteoform-spectrum-matches with at least 10 matched fragment ions. The mass graph alignment algorithm identified all the 1, 031 matches reported by MS-Align-E as well as 152 proteoform-spectrum-matches missed by MS-Align-E, of which 144 correspond to proteoforms with terminal truncation. The main reason why the identifications were missed by MS-Align-E is that MS-Align-E is not able to identify truncated proteoforms. The comparison demonstrated that the mass graph alignment method outperformed MS-Align-E in identifying truncated proteoforms.

## 4 Conclusions

In this paper, we proposed the mass graph representation of proteoforms and MS/MS spectra and transformed the proteoform identification problem to the semi-global mass graph alignment problem. In addition, we proposed dynamic programming algorithms for the RMGA problem, a restricted version of the mass graph alignment problem. The experiments on a histone H4 top-down MS/MS data set showed that the proposed mass graph alignment method is more powerful than MS-Align-E in identifying truncated proteoforms.

## Appendix A The NP-hardness of the mass graph alignment problem

In the decision version of the mass graph alignment problem, we are given a proteoform mass graph *G*, a spectral mass graph *H*, and a threshold *t*, the objective is to determine if there exist a path *A* in *G* and a path *B* in *H* such that Score(*A, B*) ≥ *t*.

### Theorem 1.

*The decision version of the mass graph alignment problem is NP-complete*.

#### Proof.

We reduce the subset sum problem [5] to the decision version of the mass graph alignment problem. Given a set *S* of positive integers and a target positive integer *b*, the subset sum problem is to determine if there is a subset *R* ⊆ *S* such that the sum of the numbers in *R* equals *b*. For a given instance *S* = {*a*_1_, *a*_2_*,…,a_n_*} of the subset sum problem, we construct an instance of the mass graph alignment problem using the following method. First, a proteoform graph *G* is constructed with two steps: (1) a total of *n*+1 nodes *x*_0_*, x*_1_*,…,x_n_* are added to the graph; (2) for each pair of neighboring nodes *x_i−_*_1_*, x_i_* (1 ≤ *i* ≤ *n*), a black directed edge and a red directed edge are added from *x_i−_*_1_ to *x_i_*. The weight for the red edge is 0, and the weight for the black edge is *a_i_*. Second, we construct a spectral mass graph that contains only a start node, an end node, and a directed edge from the start node to the end node. The weight of the edge is *b*.

→ If there is a solution *R* to the instance of the subset sum problem, we can find an alignment with a shared mass count score 2. We find a path from *x*_0_ to *x_n_* as follows: if *a_i_* ∈ *R*, then we choose the black edge to connect *x_i−_*_1_ to *x_i_;* otherwise, the red edge. As a result, the weight of the path is *b*, and the score of the alignment between the path and the full path in the spectral mass graph is 2.

← If the instance of the mass graph alignment problem has an alignment with a shared mass count score 2, then the instance of the subset sum problem has a solution. Let (*A, B*) be the two paths of the alignment, where *A* is in *G* and *B* is in *H*. Since the path *B* contains only one edge with a weight *b*, the weight of *A* is also *b*. For each black edge (*x_i−_*_1_*, x_i_*) in *A*, we add *a_i_* to the subset *R*. The resulting subset is a solution to the instance of the subset sum problem.

## Appendix B The algorithm for the local RMGA problem.

**Fig. 4.**
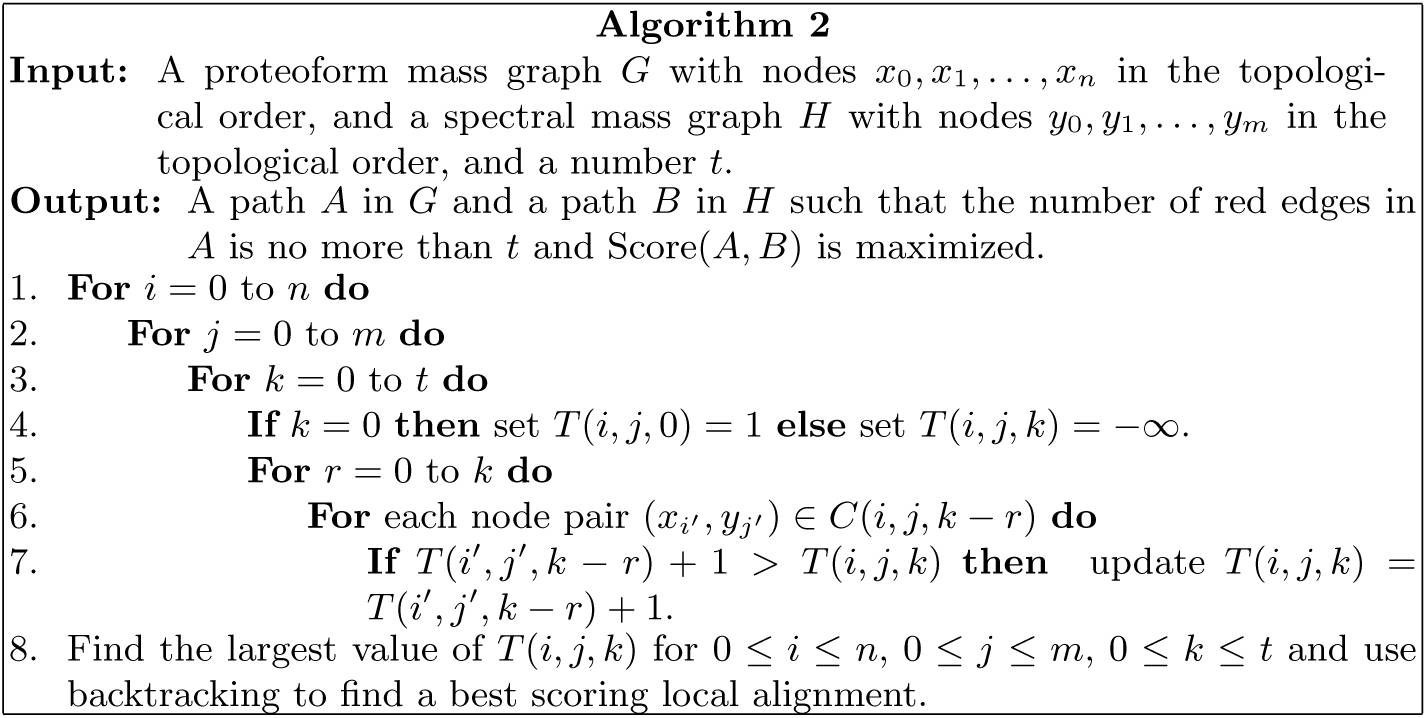
The algorithm for the local RMGA problem.

## Appendix C Variable PTMs used in the analysis of the histone data set

**Table 1.** Five variable PTMs used in the identification of proteoforms of the histone H4 protein

| PTM | Monoisotopic mass shift (Da) | Amino acids |
| --- | --- | --- |
| Acetylation | 42.01056 | R, K |
| Methylation | 14.01565 | R, K |
| Dimethylation | 28.03130 | R, K |
| Trimethylation | 42.04695 | R |
| Phosphorylation | 79.96633 | S, T, Y |

## Appendix D Histograms for the identified proteoform-spectrum-matches

**Fig. 5.**
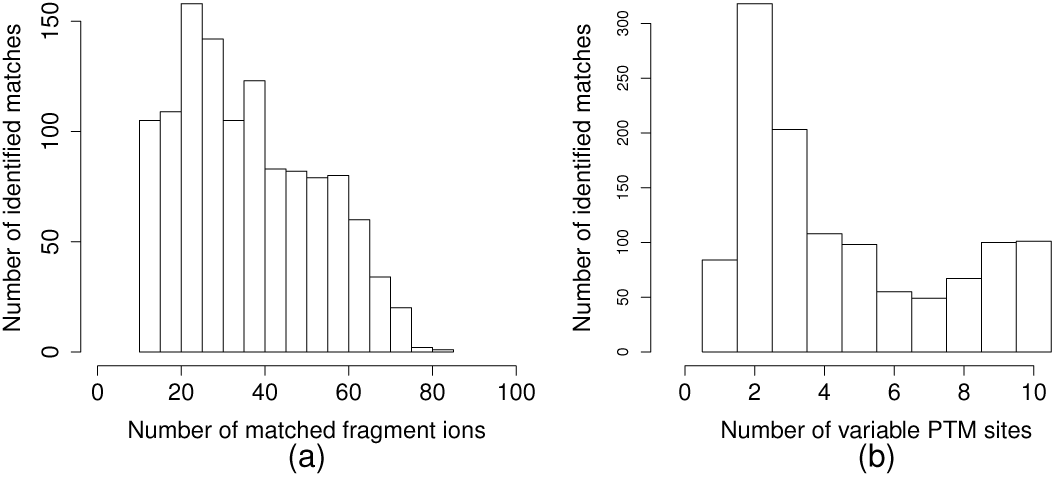
Histograms for the 1, 183 proteoform-spectrum-matches reported from the histone H4 data set by the mass graph alignment method: (a) the number of matched fragment ions; (b) the number of variable PTM sites.

